# A reductionist approach to model photosynthetic self-regulation in eukaryotes in response to light

**DOI:** 10.1101/033894

**Authors:** Anna B. Matuszyńska, Oliver Ebenhöh

## Abstract

Along with the development of several large-scale methods such as mass spectrometry or micro arrays, genome wide models became not only a possibility but an obvious tool for theoretical biologists to integrate and analyse complex biological data. Nevertheless, incorporating the dynamics of photosynthesis remains one of the major challenges while reconstructing metabolic networks of plants and other photosynthetic organisms. In this review, we aim to provide arguments that small-scale models are still a suitable choice when it comes to discover organisational principles governing the design of biological systems. We give a brief overview of recent modelling efforts in understanding the interplay between rapid, photoprotective mechanisms and the redox balance within the thylakoid membrane, discussing the applicability of a reductionist approach in modelling self-regulation in plants, and outline possible directions for further research.

---

**Abbreviations**
LHC: light harvesting complex
NPQ: non-photochemical quenching
PSI: photosystem I
PSII: photosystem II
qE: high-energy dependent quenching
qT: state transitions
RC: reaction centre
VAZ: violaxanthin-antheraxanthin-zeaxanthin cycle

## Introduction

In the process of photosynthesis solar energy is harvested by chlorophyll pigments and converted into chemical energy by a series of redox reactions. High light intensities may severely impair the photosynthetic apparatus and damage the reaction centres, where charge separation occurs. In order to protect themselves against photodamage, plants and other photosynthetic organisms are capable of switching from a photosynthetic, light-harvesting to a protective status, in which excess absorbed radiant energy is dissipated as heat [1].

Through the reorganisation of light harvesting complexes plants gain the ability to dynamically react to external stimuli and to keep the redox balance within the thylakoid membrane [2]. However, what is a desired and even essential mechanism in natural, fluctuating environments, becomes an unwanted feature in industrial cultivation, where one aims at utilising the applied light energy for photochemistry with the highest possible efficiency. Clearly, a thorough understanding of the molecular signalling mechanisms guiding acclimation responses is required to optimise biotechnological exploitation of photosynthetic organisms, for example for the production of high-value commodities. Such an understanding, which can only be obtained by combining several scientific approaches, will allow to assess, quantify and eventually minimise the rate of energy loss. In the long term, this knowledge has the potential to support increasing plant productivity, and thus contribute to solving the grand challenge of the 21st century imposed by the increasing food and energy demand [3, 4].

Theoretical approaches are powerful to discover organisational principles governing the design of biological systems. Properly constructed mathematical models verify and complement experimentally obtained results, reflect the current state of knowledge and set theoretical frameworks to derive novel hypotheses and perform investigations which are often experimentally challenging, if not impossible. Mathematical models can take many forms, depending on the research question they aim to answer [5]. By definition, models are a simplified representation of reality and can focus on different timescales and different levels of complexity (Figure 1). System-level models of metabolism found a number of applications but, because of the intrinsic assumption of a stationary state, they face the challenge of including the dynamics of photosynthesis [6] when applied to phototrophic organisms.

**Figure 1.**
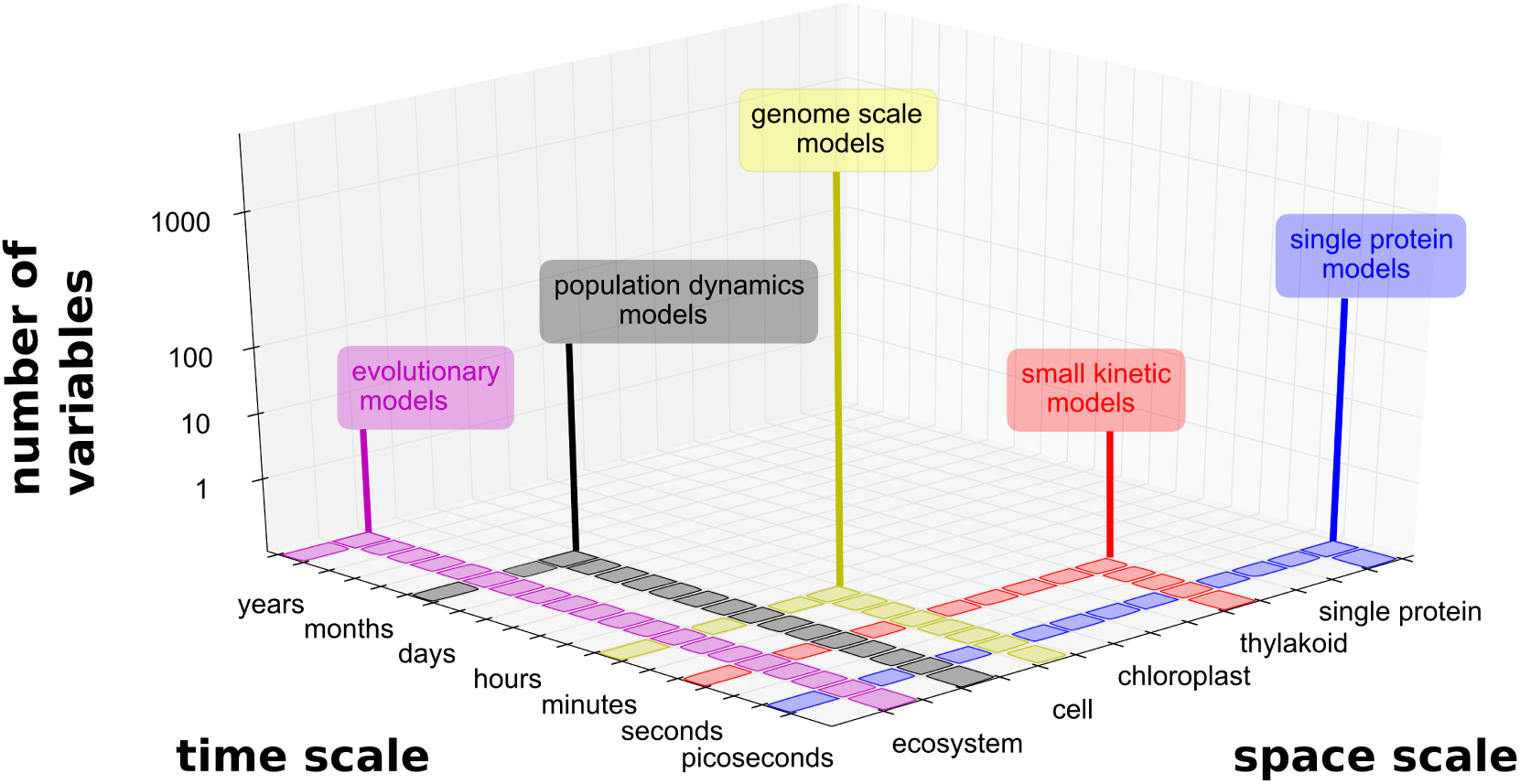
The three dimensions of model reduction. Existing photosynthetic models cover different levels of complexity and details depending on the research questions they aim to answer, ranging from the detailed models of processes occurring within PSII on the timescale of picoseconds to nanoseconds, reviewed in [7], over the biochemically structured models of culture growth in bioreactors [8, 9] to models of photosynthethic evolution [10]. All of them reduce the commplexity by focussing on selected temporal and spatial scales.

In this review, we aim to provide arguments for the application of reductionist approaches in photosynthetic research to study self regulation in plants. For that we discuss mathematical models published on the topic in the past decade and discuss challenges and future prospects associated with dynamic, differential equation-based models.

## Self regulation

In natural conditions, plants are exposed to rapid fluctuations in their environment [11] including changes in light intensity and quality. When a chlorophyll absorbs a photon it is excited to a higher energy state from which it can relax either by fuelling the photochemical reactions or by dissipating the excess energy in the form of fluorescence or heat [1]. In high light, excitation of chlorophyll may be faster than relaxation and chlorophyll singlet excited states accumulate [12]. This leads to the formation of chlorophyll triplets that, by reacting with molecular oxygen, result in highly reactive and dangerous singlet oxygen [13]. Through processes collectively named as non-photochemical quenching (NPQ) almost all eukaryotic autotrophs [14] avoid and minimise such photooxidative stress. Photosynthesis is driven by energy collected by complexes associated with two photosystems, which are preferentially excited by different wavelengths. Acclimation to fluctuating environments by balancing excitation of these two photosystems is achieved by an additional mechanism, in which light-harvesting complexes are relocated [15].

Although a number of genes and proteins involved in these acclimation pathways have been identified, in many cases the molecular basis for their dynamics remains unknown. Thus, a number of mathematical models have been developed with the goal to understand the regulatory principles and to support the identification of the underlying molecular mechanisms.

## Models

Much of today’s knowledge about the dynamics of photosynthesis was brought by reductionist models, dating back to the extremely simplified, but illustrative pioneering model of leaf photosynthesis by Thornley [16]. The question is whether this approach is still justified in the era of quantitative biology. The recent rapid advance in experimental techniques, such as mass-spectrometry and high-throughput sequencing, allows obtaining global snapshots of the status of a cell with unprecedented precision [17]. This wealth of information allows for example the reconstruction of genome-scale metabolic networks encompassing the entirety of all known biochemical reactions [18]. One would expect that with this richness of available data, a fundamental biochemical process like photosynthesis would be already well understood. In fact, a few attempts have been made to apply genome-scale metabolic models to photosynthetic organisms including plants [19], green algae [20] and cyanobacteria [21] (recently reviewed in [6]) and these approaches were successful in providing valuable insight into the dependence of stationary flux distributions to external conditions. However, the inherent steady-state assumptions in the mathematical analysis techniques [22] makes them unsuitable to explore the regulatory mechanisms underlying the dynamic responses, which are so essential for organisms that need to cope with changing environmental conditions [23].

In contrast, small-scale kinetic models are designed for an in-depth investigation of individual biological components and can provide information on the dynamics of the system, far away from the steady state, and predict temporal responses to different perturbations. Here, reduced model complexity and low numbers of model parameters support the process of creating a general theoretical platform to study mechanisms conserved over a wide range of species, including the plant kingdom, but also other photosynthetic microalgae. In the past decade a handful of new kinetic models have been published with the aim to help understand underlying principles governing short term acclimation mechanisms. Due the fact that the effect of regulatory acclimation mechanisms can be easily monitored in a minimally invasive way by chlorophyll fluorescence measurements, many of the existing models aim at simulating the dynamics of the fluorescence signal [24, 25].

### Models of qE

The major and most rapid component of NPQ, termed energy-dependent quenching (qE) [1], relaxes within seconds to minutes and is triggered by a high proton gradient (ΔpH) over the thylakoid membrane [28]. General consensus is that both photochemical and non-photochemical quenching are mainly associated with light harvesting complexes of photosystem II [29] and therefore models investigating qE are commonly reduced to include only the essential reactions around PSII and focus on depicting the chlorophyll fluorescence kinetics. Recently, a simplistic three-state model of the reaction centres [30] was complemented with a data-derived, heuristic sigmoid function of qE-quenching activity [31]. This approach allowed for quantitative predictions of the state of the photosynthetic apparatus under varying light conditions while expanding the parameters set moderately to only 13 parameters. Nevertheless it does not provide novel mechanistic explanations of the regulation of heat dissipation. To understand the precise role of the known quenching components, more mechanistic models are needed.

In [27], a minimal mathematical model of NPQ is presented, which reduces the system to only three differential equations and the system boundary is drawn at the cytochrome b_6_f complex. The qE mechanism is simplified to include only one pH dependent component, ignoring for example details on the dynamics and the precise role of the xanthophyll cycle. In [32], a more detailed and accurate model of quenching is presented, however at the cost of drastically increased complexity (26 non-linear differential equations). Here, the rate of qE activity depends on two components, protonation of the PsbS protein and operation of the violaxanthin, antheraxanthin, zeaxanthin (VAZ) cycle. The model analysis suggested that, despite its pH-dependency, qE does not affect the lumen pH in plants and therefore does not regulate the mode of electron flow [32].

Existing mathematical models of high-energy state quenching are able to reproduce the main biological features of quenching activity [27] and help to quantify the beneficial impact of qE under fluctuating light conditions [32]. A review of mathematical models and measurements of energy-dependent quenching was published in [33].

### Models of state transitions

An alternative mechanism to reduce the amount of excitation energy is by decreasing the delivery rate of photons to the PSII reaction centre (RC). In the process termed *state transitions* (qT), major light harvesting complexes (LHCs) that are usually associated with PSII separate from the photosystem and move towards PSI, balancing the overall excitation and regulating the production of ATP [34], but not necesarly switching between cyclic and linear electron flow [35] as previously reported [36]. The development of mathematical models of state transitions is challenging due to the limited information regarding the exact molecular mechanisms governing this process. State transitions are triggered by the imbalance in the redox poise of the PQ pool, where an overreduced pool indirectly activates a kinase [37] (STN7 in *A. thaliana*, Stt7 in *C. reinhardtii)*, that phosphorylates antenna associated with PSII, triggering reversible antenna movement, thus resulting in a decreased delivery of photons to the PSII RC, therefore reducing PSII activation [38]. To our knowledge, the only available dynamic model of state transitions has been published recently by one of the authors [39]. This model provides a reliable representation of state transitions and presents a good basis for analysing the entire photosynthetic electron transport chain and its interaction with environmental cues and downstream processes. Further, an attempt was made to link both quenching components qE and qT into a single model. The discrepancies between simulated and experimentally obtained fluorescence traces suggested that the conventional view, where all phosphorylated antennae that detach from PSII will become associated with PSI, might be too simplistic. Indeed, a new concept of free antennae was recently introduced [40, 41] revisiting a well established view on the role of state transitions. These new experimental findings open new perspectives for modellers to underpin novel proposed mechanisms with theoretical studies.

### Models of both mechanisms

Our current understanding of dynamic regulations of photosynthesis to light variations indicate the existence of a complex regulatory network [15]. To fully understand the principles according to which this network operates (Figure 2A) and to which degree it is conserved among photosynthetic organisms, we require a model that includes both mechanisms. Considerations of only PSII reactions for qE studies (Figure 2B) or only the PQ balance for qT investigations (Figure 2C) are not sufficient, because both mechanisms are not operating independently and are likely to affect each other (Figure 2D).

**Figure 2.**
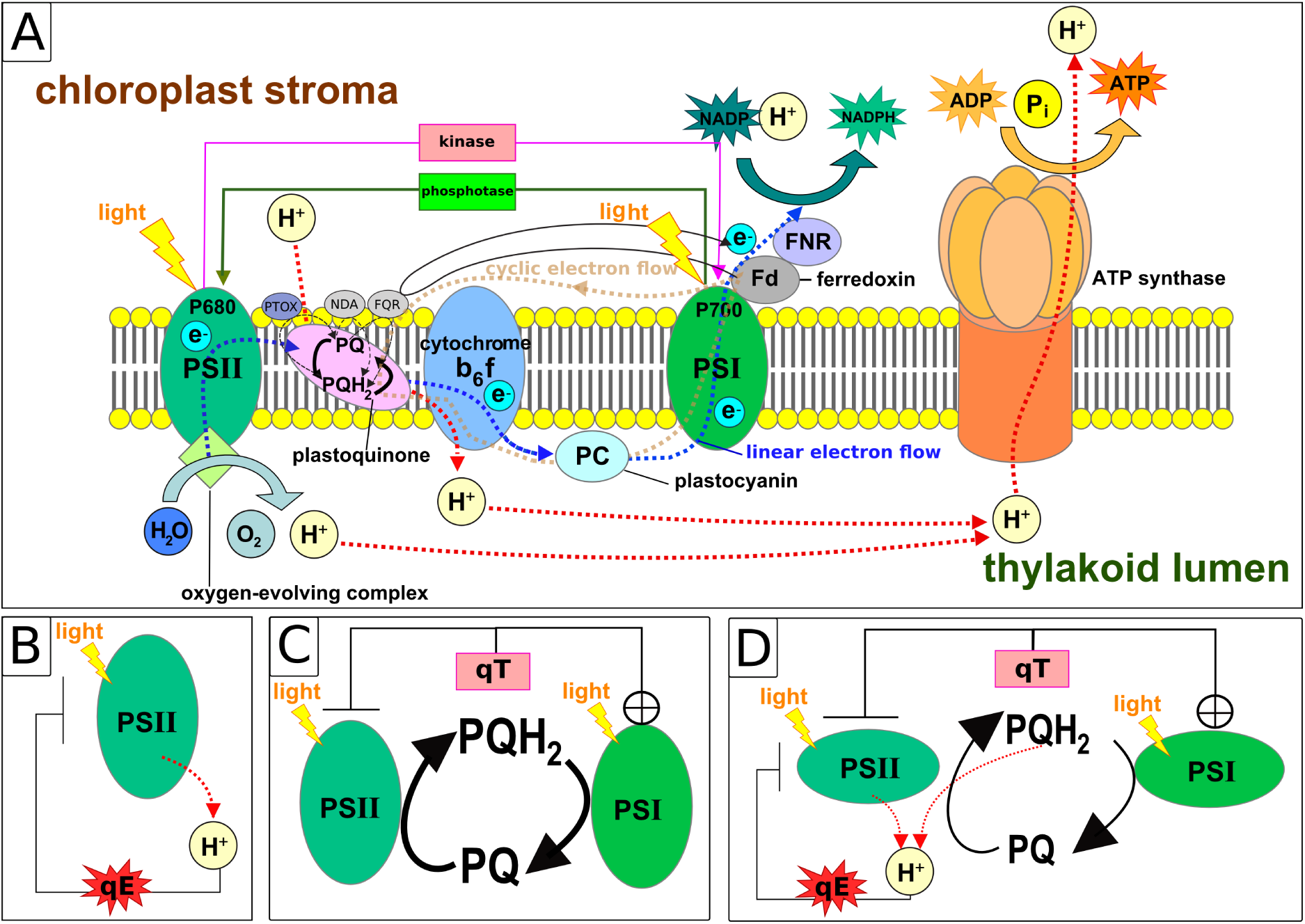
Illustration of reductionist approaches in modelling photosynthetic acclimation mechanisms. (A) Schematic representation of the photosynthetic electron transfer chain with a high level of complexity including both linear and cyclic electron flow and movement of antennae governed by the kinase-phosphotase pair. Figure modified from [26] (B) Based on our current understanding of qE, we can reduce the system to include only reactions within PSII [27] and simplify the quenching mechanism as a direct consequence of the acidification of the thylakoid lumen. (C) Similarly, to study state transitions we can reduce the system to two photosystems, which antagonistically reduce and oxidise the plastoquinone pool. A reduced pool will activate a kinase, triggering relocation of antennae from PSII to PSI, thus inhibiting PSII activity and activating PSI. (D) The two mechanisms are not independent, therefore a model that includes both mechanisms is required to reveal the principles of photosynthetic self-regulation under different light conditions.

So far, to the best of our knowledge, there is no such unifying model for photosynthetic eukaryotes available. Even the most comprehensive dynamic mechanistic model of C3 photosynthesis published to date [42] does not account for state transitions. We have therefore applied the reductionist approach to construct a minimal model including state transitions as described in [39], the mechanisms of energy-dependent quenching as described in [32], while aiming for a reduced complexity in the spirit of the minimal model in [27].

Preliminary results of this combined model are already highly illustrative and are able to explain why two regulatory mechanisms are needed and in which light regime each mechanism dominates. In Figure 3, the steady-state redox state of the plastoquinone pool is depicted for an *in silico* experiment, which is difficult to realise experimentally: In the absence of state transitions, the total light intensity (*x*-axis) and the fraction of light absorbed by PSII (*y*-axis) were varied. It can be observed that for low light intensities the redox state of the PQ pool exhibits a sharp transition from oxidised to reduced for an increasing percentage of light absorbed by PSII.

**Figure 3.**
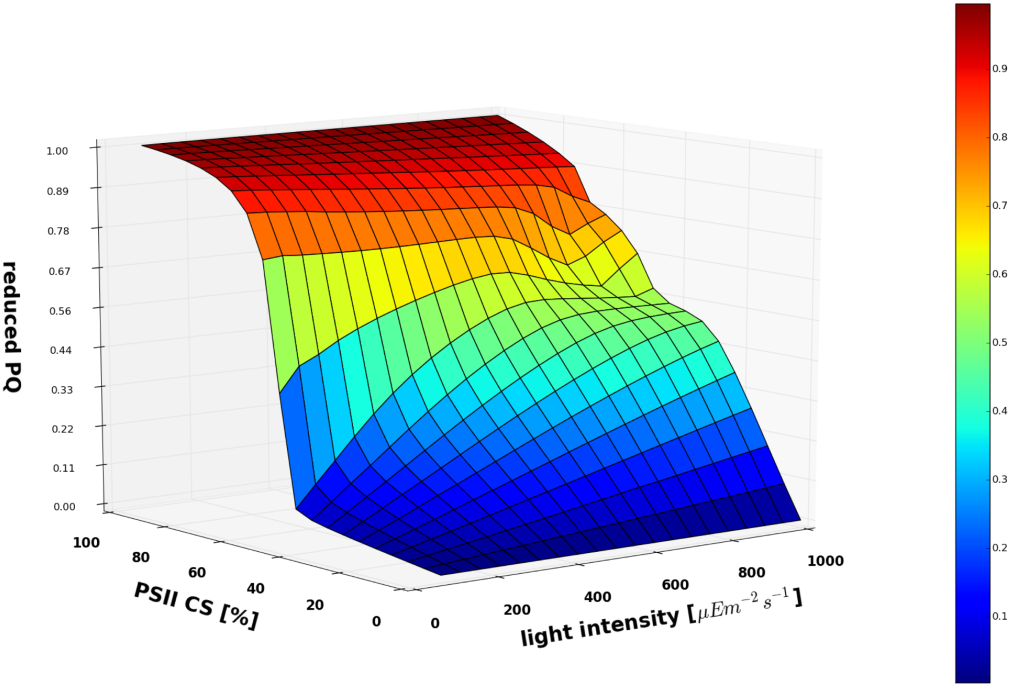
Steady state analysis. Steady state redox state of the plastoquinone pool for a model in which state transitions are not effective reveals the importance of this mechanism under low light conditions. The predicted steady state of the redox state of the PQ pool is plotted in dependency on the total light intensity and the fraction of energy absorbed by PSII.

This demonstrates the importance of the capacity to regulate the relative energy transfer to PSI and PSII for the maintenance of a healthy redox balance required for an efficient photosynthetic electron transport. For higher light intensities the transition becomes increasingly more gradual. The energy dependent dissipation of absorbed energy as heat (qE) leads to an increasingly pronounced plateau-like behaviour, illustrating the importance of qE under high light conditions.

## Future directions

Evidently, there are still many unanswered questions and reaching a true understanding of the photosynthetic self-regulatory system still requires considerable effort. In our opinion, a dynamic model that encompasses both major shortterm acclimation mechanisms has the potential to address central unresolved problems and thus will help to develop encompassing theoretical concepts further. In particular, we see a need to address a number of key areas.

*Photosynthetic mutants* are widely used to study photoregulatory mechanisms in plants and green algae. In order to draw from the multitude of genetic and phenotypic information available and incorporate this information into the theory building process, a model is required, which incorporates more than one regulatory mechanism. For example, the theoretical analysis of the metabolic signals in the *C. reinhardtii* double mutant *npq4stt7-9*, which is deficient in state transitions and impaired in expression of a protein required for qE activation [43], can only be performed with a model describing both qT and qE. The urgency to develop such a model is further stressed by the observation that removing one photoprotective mechanisms triggers compensatory responses, where the remaining mechanism partly takes over the role of the removed one [43].

It has been demonstrated that plants possess a *cellular light memory*, where they can physiologically memorise previous light exposure periods to improve their light acclimatory and immune defence responses even days after exposure [44]. To test if there exists a short-term light memory that improves environmental fitness in the time-scale of minutes to hours, we expanded the minimal mathematical model presented in [27] by including the slower component of NPQ and mechanisms responsible for the regularly observed transient quenching induction under low light conditions [45]. Our working hypothesis states that short-term light memory is established by the slow quenching component (usually attributed to the accumulation of zeaxanthin) and the simulations of this unpublished model indeed seem to support this notion (Figure 4).

**Figure 4.**
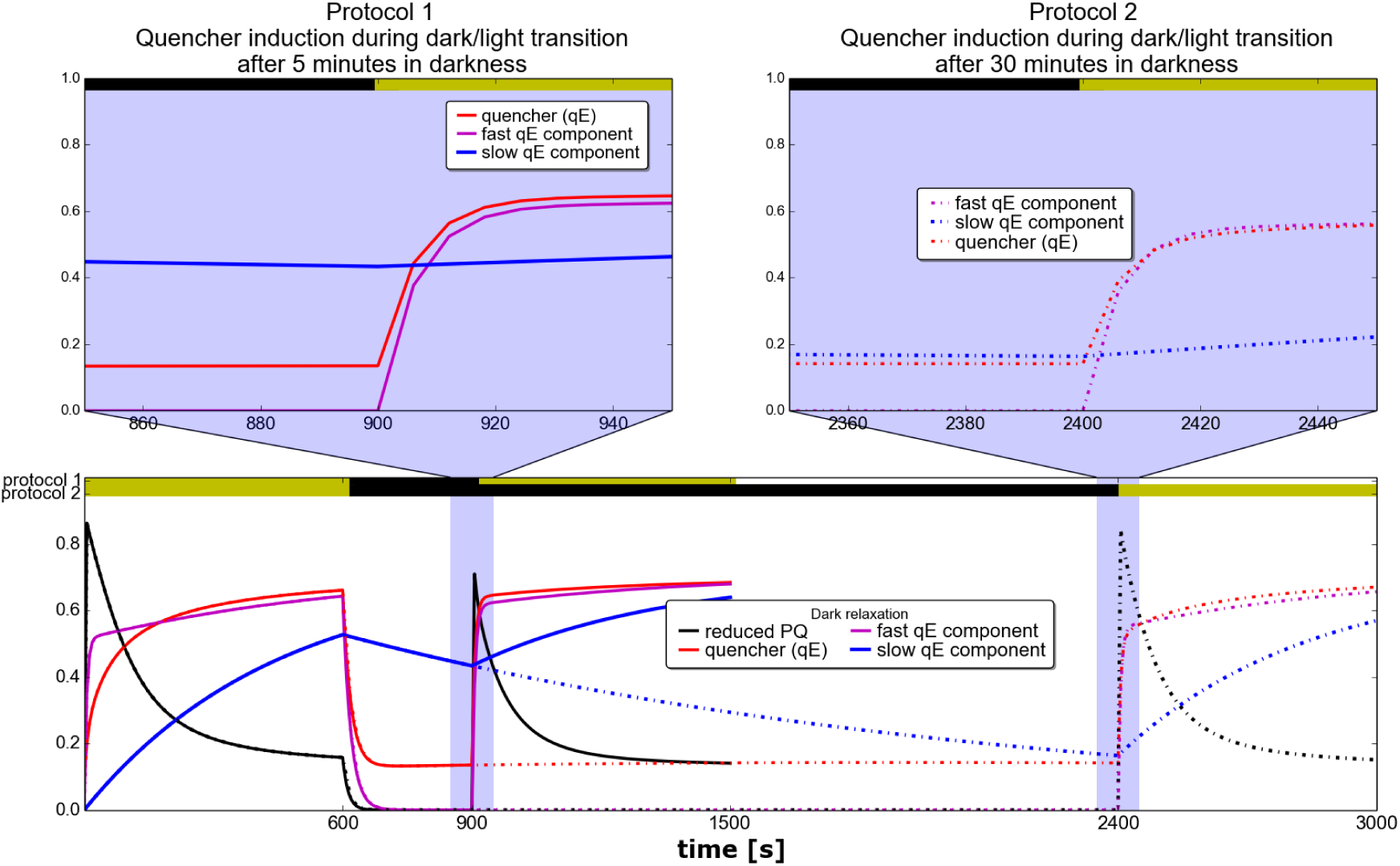
Simulations of a short-term light memory. Simulated response of Arabidopsis exposed to white light of 500 *μEm^−2^s^−1^* for 10 minutes, shifted to darkness for 5 minutes (protocol 1) or 30 minutes (protocol 2, dotted lines), in which the quenching components relax, and subsequent re-exposure to light of the original intensity. Simulations were performed in Python using a kinetic model developed from [27], complemented by the slower component of NPQ and by mechanisms responsible for the transiently generated NPQ under low light conditions.

It is widely accepted that *light quality* is another major trigger of regulatory responses after light intensity [46]. However, how exactly different light frequencies affect the extent and rate of NPQ is largely unknown. First approaches to implement a spectral dependency of the light reactions were presented in [20] in the context of the genome-scale metabolic network reconstruction of *C. reinhardtii*, where a prism reaction was added as an intermediate step between incident and absorbed light. This approach led to a better understanding of algae growth under different spectral compositions, but neither can it help to understand the dynamics of the regulation nor can it provide detailed information about the energy loss due to photoprotective mechanisms. We suggest that these questions can be best addressed by modifying small kinetic models to include a description of a frequency-dependent excitation of the chlorophylls and reaction centres.

Last but not least, we are convinced that the selective reduction of certain processes can help to identify common underlying principles of NPQ in different species. For example, reducing the xanthophyll cycle to its essential feature, namely that a xanthophyll can be de-epoxidised in a pH-dependent manner and epoxidised in a pH-independent way [47], allows to include species as different from plants as diatoms into the model description, where the xanthophyll cycle operates according to the same principles as in higher plants and green algae, but the molecular nature of the xanthophylls is very different [14].

## Discussion

A major challenge in mathematical model development remains the acquisition of accurate numerical values of the numerous photosynthetic parameters that are not trivial to measure *in vivo*. This is particularly pronounced while developing unifying modelling frameworks valid for different species. For that, efforts taken by Antal *et al*. [48] serve as an invaluable comprehensive data base and will greatly help to overcome this bottleneck of model development. Hovewer, no matter how many parameters are measured, there will always be the need to fit the remaining ones to experimental curves. However, this fitting procedure can be considerably facilitated by a simple model structure, and the smaller the number of parameters, the lower the risk of overfitting. At the same time, models can provide quantitative information about values where direct measurements *in vivo* are not possible, as was for example performed for the prediction of thylakoid lumenal pH [49].

In conclusion, a number of existing mathematical models can capture the fluorescence dynamics under changing light conditions and support the understanding of photosynthetic self-regulatory mechanisms, which could not be obtained by other modelling or experimental approaches. Nevertheless, to reach a more complete understanding of photosynthesis and its regulation, it will become necessary to incorporate dynamic regulatory models with genome-scale approaches. In order to reach this stage, we need to understand how to upscale the detailed insight into regulatory dynamics and how to capture the essential features in a way compatible with large-scale model descriptions. To reach this understanding, we see the necessity to further pursue the reductionist approach resulting in small kinetic models, because only these will facilitate the formulation of unifying theoretical frameworks and discover the significance of the individual regulatory mechanisms for photosynthetic efficiency.

## Financing

This work was supported by the Marie Curie Initial Training Network AccliPhot financed by the European Union [grant number PITN-GA-2012-316427 (to A.M. and O.E.)]; and the Deutsche Forschungsgemeinschaft [Cluster of Excellence on Plant Sciences, CEPLAS (EXC 1028) (to O.E.)].

## References

[1] P. Müller, X.-P. Li, and K. K. Niyogi, “Nonphotochemical quenching. a response to excess light energy.,” Plant Physiol, vol. 125, pp. 1558–1566, Apr 2001.

[2] J. Minagawa, “Dynamic reorganization of photosynthetic supercomplexes during environmental acclimation of photosynthesis.,” Front Plant Sci, vol. 4, p. 513, 2013.

[3] N. Powell, X. Ji, R. Ravash, J. Edlington, and R. Dolferus, “Yield stability for cereals in a changing climate,” Funct. Plant Biol., vol. 39, no. 7, pp. 539–552, 2012.

[4] N. P. A. Hüner, K. Dahal, L. V. Kurepin, L. Savitch, J. Singh, A. G. Ivanov, K. Kane, and F. Sarhan, “Potential for increased photosynthetic performance and crop productivity in response to climate change: role of CBFs and gibberellic acid.,” Front. Chem., vol. 2, no. April, p. 18, 2014.

[5] T. Pfau, N. Christian, and O. Ebenhöh, “Systems approaches to modelling pathways and networks,” Brief. Funct. Genomics, vol. 10, no. 5, pp. 266–279, 2011.

[6] C. Baroukh, R. Muñoz Tamayo, J.-P. Steyer, and O. Bernard, “A state of the art of metabolic networks of unicellular microalgae and cyanobacteria for biofuel production,” Metab. Eng., vol. 30, pp. 49–60, 2015.

[7] D. Lazár and J. Jablonský, “On the approaches applied in formulation of a kinetic model of photosystem ii: Different approaches lead to different simulations of the chlorophyll a fluorescence transients,” Journal of Theoretical Biology, vol. 257, no. 2, pp. 260 – 269, 2009.

[8] J.-F. Cornet, C. Dussap, and J.-B. Gros, “Kinetics and energetics of photosynthetic micro-organisms in photobioreactors,” in Bioprocess and Algae Reactor Technology, Apoptosis, vol. 59 of *Advances in Biochemical Engineering Biotechnology*, pp. 153–224, Springer Berlin Heidelberg, 1998.

[9] G. Cogne, M. Rügen, A. Bockmayr, M. Titica, C. G. Dussap, J. F. Cornet, and J. Legrand, “A model-based method for investigating bioenergetic processes in autotrophically growing eukaryotic microalgae: Application to the green algae Chlamydomonas reinhardtii,” Biotechnol. Prog., vol. 27, no. 3, pp. 631–640, 2011.

[10] D. Heckmann, S. Schulze, A. Denton, U. Gowik, P. Westhoff, A. P. M. Weber, and M. J. Lercher, “XPredicting C4 photosynthesis evolution: Modular, individually adaptive steps on a mount fuji fitness landscape,” Cell, vol. 153, no. 7, 2013.

[11] E. Kaiser, A. Morales, J. Harbinson, J. Kromdijk, E. Heuvelink, and L. F. M. Marcelis, “Dynamic photosynthesis in different environmental conditions,” J. Exp. Bot., pp. 1–12, 2014.

[12] G. Bartosz, “Oxidative stress in plants,” Acta Physiol. Plant., vol. 19, no. 1, pp. 47–64, 1997.

[13] M. Ballottari, M. Mozzo, J. Girardon, R. Hienerwadel, and R. Bassi, “Chlorophyll triplet quenching and photoprotection in the higher plant monomeric antenna protein Lhcb5,” J. Phys. Chem. B, vol. 117, no. 38, pp. 11337–11348,2013.

[14] R. Goss and B. Lepetit, “Biodiversity of NPQ.,” J. Plant Physiol., 2014.

[15] J. Minagawa and R. Tokutsu, “Dynamic regulation of photosynthesis in chlamydomonas reinhardtii,” The Plant Journal, vol. 82, no. 3, pp. 413–428, 2015.

[16] J. H. M. Thornley, “Light fluctuations and photosynthesis,” Ann Bot, vol. 38, pp. 363–373, 1974.

[17] F. D. Mast, A. V. Ratushny, and J. D. Aitchison, “Systems cell biology.,” J Cell Biol, vol. 206, pp. 695–706, Sep 2014.

[18] D. A. Fell, M. G. Poolman, and A. Gevorgyan, “Building and analysing genome-scale metabolic models.,” Biochem Soc Trans, vol. 38, pp. 1197–1201, Oct 2010.

[19] M. G. Poolman, M. G. Poolman, L. Miguet, L. Miguet, L. J. Sweetlove, L. J. Sweetlove, D. A. Fell, and D. A. Fell, “A Genome-scale Metabolic Model of Arabidopsis thaliana and Some of its Properties.,” Plant Physiol., vol. 151, no. November, pp. 1570–1581, 2009.

[20] R. L. Chang, L. Ghamsari, A. Manichaikul, E. F. Y. Hom, S. Balaji, W. Fu, Y. Shen, T. Hao, B. Ø. Palsson, K. Salehi-Ashtiani, and J. A. Papin, “Metabolic network reconstruction of chlamydomonas offers insight into light-driven algal metabolism.,” Mol Syst Biol, vol. 7, p. 518, 2011.

[21] H. Knoop, Y. Zilliges, W. Lockau, and R. Steuer, “The metabolic network of synechocystis sp. pcc 6803: systemic properties of autotrophic growth.,” Plant Physiol, vol. 154, pp. 410–422, Sep 2010.

[22] K. J. Kauffman, P. Prakash, and J. S. Edwards, “Advances in flux balance analysis.,” Curr Opin Biotechnol, vol. 14, pp. 491–496, Oct 2003.

[23] E. Collakova, J. Y. Yen, and R. S. Senger, “Are we ready for genome-scale modeling in plants?,” Plant Sci., vol. 191–192, pp. 53–70, 2012.

[24] K. Maxwell and G. Johnson, “Chlorophyll fluorescence,” J. Exp. Bot., vol. 51, no. 345, pp. 659–668, 2000.

[25] A. Stirbet, G. Y. Riznichenko, A. B. Rubin, and Govindjee, “Modeling chlorophyll a fluorescence transient: relation to photosynthesis.,” Biochem. Biokhimiya, vol. 79, no. 4, pp. 291–323, 2014.

[26] WikimediaCommons, “Light-dependent reactions of photosynthesis in the thylakoid membrane of plant cells. Redrawn and formatted for better quality SVG.,” 2015. File:Thylakoid_membrane_3.svg.

[27] O. Ebenhöh, T. Houwaart, H. Lokstein, S. Schlede, and K. Tirok, “A minimal mathematical model of nonphotochemical quenching of chlorophyll fluorescence.,” Biosystems., vol. 103, pp. 196–204, Feb. 2011.

[28] C. A. Wraight and A. R. Crofts, “Delayed fluorescence and the high-energy state of chloroplasts.,” Eur. J. Biochem., vol. 19, no. 3, pp. 386–397, 1971.

[29] P. Horton, a. V. Ruban, and R. G. Walters, “Regulation of Light Harvesting in Green Plants,” Annu. Rev. Plant Physiol. Plant Mol. Biol., vol. 47, no. 1, pp. 655–684, 1996.

[30] B.-P. Han, “A mechanistic model of algal photoinhibition induced by photodamage to photosystem-II.,” Journal of theoretical biology, vol. 214, pp. 519–527, 2002.

[31] A. Nikolaou, A. Bernardi, A. Meneghesso, F. Bezzo, T. Morosinotto, and B. Chachuat, “A model of chlorophyll fluorescence in microalgae integrating photoproduction, photoinhibition and photoregulation.,” Journal of biotechnology, vol. 194, pp. 91–9, Jan. 2015.

[32] J. Zaks, K. Amarnath, D. M. Kramer, K. K. Niyogi, and G. R. Fleming, “A kinetic model of rapidly reversible nonphotochemical quenching.,” Proc. Natl. Acad. Sci. U. S. A., vol. 109, pp. 15757–62, Sept. 2012.

[33] J. Zaks, K. Amarnath, E. J. Sylak-Glassman, and G. R. Fleming, “Models and measurements of energy-dependent quenching.,” Photosynth. Res., vol. 116, pp. 389–409, Oct. 2013.

[34] J.-D. Rochaix, “Regulation and dynamics of the light-harvesting system.,” Annual review of plant biology, vol. 65, pp. 287–309, Jan. 2014.

[35] H. Takahashi, S. Clowez, F.-A. Wollman, O. Vallon, and F. Rappaport, “Cyclic electron flow is redox-controlled but independent of state transition.,” Nature communications, vol. 4, p. 1954, Jan. 2013.

[36] G. Finazzi, A. Furia, R. P. Barbagallo, and G. Forti, “State transitions, cyclic and linear electron transport and photophosphorylation in Chlamydomonas reinhardtii,” Biochim. Biophys. Acta - Bioenerg., vol. 1413, no. 3, pp. 117–129, 1999.

[37] F. Zito, G. Finazzi, R. Delosme, W. Nitschke, D. Picot, and F. A. Wollman, “The Qo site of cytochrome b6f complexes controls the activation of the LHCII kinase,” EMBO J., vol. 18, no. 11, pp. 2961–2969, 1999.

[38] J.-D. Rochaix, “Reprint of: Regulation of photosynthetic electron transport.,” Biochim Biophys Acta, vol. 1807, pp. 878–886, Aug 2011.

[39] O. Ebenhöh, G. Fucile, G. Finazzi, J.-D. Rochaix, and M. Goldschmidt-Clermont, “Short-term acclimation of the photosynthetic electron transfer chain to changing light: a mathematical model,” Philosophical Transactions of the Royal Society B: Biological Sciences, vol. 369, no. 1640, 2014.

[40] G. Nagy, R. Ünnep, O. Zsiros, R. Tokutsu, K. Takizawa, L. Porcar, L. Moyet, D. Petroutsos, G. Garab, G. Finazzi, and J. Minagawa, “Chloroplast remodeling during state transitions in Chlamydomonas reinhardtii as revealed by noninvasive techniques in vivo.,” Proc. Natl. Acad. Sci. U. S. A., vol. 111, pp. 5042–7, Apr. 2014.

[41] C. Unlü, B. Drop, R. Croce, H. van Amerongen, H. V. Amerongen, and C. Ünlü, “State transitions in Chlamy-domonas reinhardtii strongly modulate the functional size of photosystem II but not of photosystem I.,” Proc. Natl. Acad. Sci. U. S. A., vol. 111, pp. 3460–5, 2014.

[42] X.-G. Zhu, Y. Wang, D. R. Ort, and S. P. Long, “e-photosynthesis: a comprehensive dynamic mechanistic model of C3 photosynthesis: from light capture to sucrose synthesis.,” Plant. Cell Environ., vol. 36, pp. 1711–27, Sept. 2013.

[43] G. Allorent, R. Tokutsu, T. Roach, G. Peers, P. Cardol, J. Girard-Bascou, D. Seigneurin-Berny, D. Petroutsos, M. Kuntz, C. Breyton, F. Franck, F.-A. Wollman, K. K. Niyogi, A. Krieger-Liszkay, J. Minagawa, and G. Finazzi, “A dual strategy to cope with high light in Chlamy-domonas reinhardtii.,” Plant Cell, vol. 25, pp. 545–57, Feb. 2013.

[44] M. Szechyńska-Hebda, J. Kruk, M. Górecka, B. Karpińska, and S. Karpiński, “Evidence for light wavelength-specific photoelectrophysiological signaling and memory of excess light episodes in Arabidopsis.,” Plant Cell, vol. 22, pp. 2201–18, July 2010.

[45] G. Finazzi, G. N. Johnson, L. Dall’Osto, P. Joliot, F.-A. Wollman, and R. Bassi, “A zeaxanthin-independent nonphotochemical quenching mechanism localized in the photosystem II core complex.,” Proc. Natl. Acad. Sci. U. S. A., vol. 101, no. 33, pp. 12375–12380, 2004.

[46] M. J. Dring, “Photocontrol of Development in Algae,” Ann Rev, Ann Physiol, Plant Mol Biol, vol. 39, pp. 157–74, 1988.

[47] P. Jahns and A. R. Holzwarth, “The role of the xanthophyll cycle and of lutein in photoprotection of photosystem II,” Biochim. Biophys. Acta - Bioenerg., vol. 1817, no. 1, pp. 182–193, 2012.

[48] T. K. Antal, I. B. Kovalenko, A. B. Rubin, and E. Tyystjärvi, “Photosynthesis-related quantities for education and modeling,” Photosynth. Res., vol. 117, no. 1–3, pp. 1–30, 2013.

[49] D. Kramer, C. Sacksteder, and J. Cruz, “How acidic is the lumen?,” Photosynthesis Research, vol. 60, no. 2–3, pp. 151–163, 1999.

